# Deoxycholic acid (DCA) alleviates LPS-induced Inflammatory Bone Loss via modulating the “Gut-Bone” homeostasis

**DOI:** 10.1101/2025.09.26.678724

**Authors:** Sumedha Yadav, Swati Rajput, Chaman Saini, Megha Sharma, Pradyumna K. Mishra, Rupesh K. Srivastava

**Affiliations:** Translational Immunology, Osteoimmunology & Immunoporosis Lab (TIOIL), An ICMR Collaborating Centre for Excellence on Bone Health, Department of Biotechnology, All India Institute of Medical Sciences (AIIMS), New Delhi-110029, India; Department of Molecular Biology, ICMR-National Institute for Research in Environmental Health, Bhopal, MP, 462001, India

**Keywords:** Osteoporosis, Deoxycholic acid, Gut microbiota, Gut-associated metabolites, Inflammatory bone loss, Lipopolysaccharide

## Abstract

Osteoporosis and other forms of inflammatory bone loss are marked by disrupted bone remodelling resulting from an imbalance between osteoclast-mediated bone resorption and osteoblast-driven bone formation. This imbalance is often exacerbated by chronic inflammation and gut microbiota dysbiosis. Recently, attention has turned to gut-associated metabolites (GAMs) such as secondary bile acids, particularly deoxycholic acid (DCA), which act as immunomodulators influencing both systemic inflammation and bone metabolism. In this study, we investigated the role of DCA in inflammatory bone loss using an *in vivo* model, with a focus on osteoblast and osteoclast function, gut barrier integrity, and gut-microbiota diversity. DCA administration significantly suppressed osteoclastogenesis along with enhancing osteoblastogenesis, indicating its dual regulatory role in bone remodelling. Furthermore, DCA treatment enhanced gut integrity, reversed dysbiosis and reduced systemic inflammation by downregulating osteoclastogenic cytokines (TNF-α, IL-6, IL-17, RANKL, etc.). These findings suggest that DCA mitigates LPS-induced inflammatory bone loss through a multifaceted mechanism involving direct effects on bone cells and restoration of gut integrity and homeostasis. Our results highlight the therapeutic potential of targeting the gut microbiota-derived bile acid pathway, particularly DCA, as a novel strategy for managing osteoporosis and other inflammatory bone disorders.

Figure 9.
Graphical Abstract.
Secondary bile acids act via FXR and TGR5 receptors on osteoclast and osteoblast cells and thereby regulate bone remodelling.

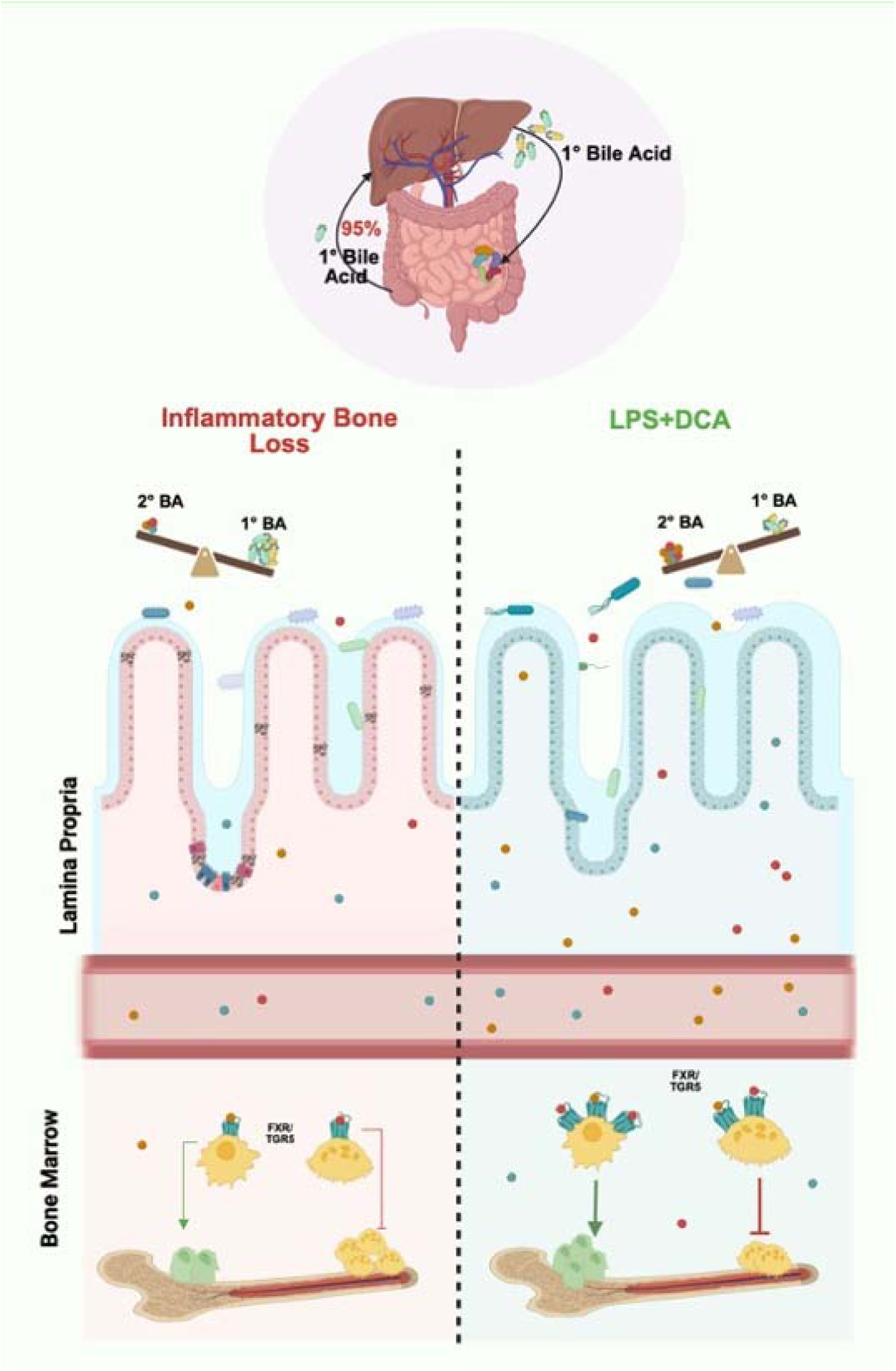

## 1.0 Introduction

Osteoporosis is a long-term bone disease characterised by decreased bone mass and the progressive breakdown of bone structure, leading to an increased susceptibility to fractures from minimal trauma or stress. It is estimated that more than 500 million people worldwide suffer from osteoporotic fractures, which leads to increased risk of morbidity, mortality, and a huge socio-economic burden [1].

Previous studies have primarily focused on the endocrine pathways involved in osteoporosis; however, recent evidence highlights the significant contribution of inflammation to its pathogenesis [2,3]. Bone is a highly dynamic organ that undergoes continuous remodelling through the coordinated action of osteoclasts, which mediate bone resorption and osteoblasts, which drive bone formation. Under physiological conditions, bone formation and resorption are tightly balanced; however, during inflammation, this equilibrium is disrupted, favouring excessive resorption, resulting in bone loss. Inflammatory markers such as interleukin-6 (IL-6), IL-17, and tumour necrosis factor alpha (TNF-α) are known to be associated with fracture risk [4]. Furthermore, chronic inflammatory conditions such as rheumatoid arthritis (RA)[5], inflammatory bowel disease (IBD)[6], diabetes [7], and pancreatitis[8] are strongly associated with reduced bone mineral density (BMD). Mobility limitation, nutritional malabsorption, and hormonal changes are further contributing factors, in addition to the related underlying inflammatory pathophysiology, that may increase the likelihood of developing secondary osteoporosis in the above-mentioned disorders.

Lipopolysaccharide (LPS) is a major constituent of the outer membrane of gram-negative bacteria, which triggers the production of pro-inflammatory cytokines such as interleukin-1β (IL-1β) and TNF-α, as well as other inflammatory mediators [9]. Receptor Activator of Nuclear Factor-kappa B Ligand (RANKL) expression is increased by this cytokine surge, which in turn stimulates osteoclastogenesis to increase bone resorption and inhibits osteoblastogenesis to limit bone growth. [10]. Therefore, a potential therapeutic approach for managing inflammatory bone loss disorders is the use of medications that specifically target LPS.

The gut-immune-bone axis has drawn a lot of attention lately. Approximately 100 trillion microorganisms inhabit the human body, with the majority residing in the gastrointestinal tract. Studies have observed that gut dysbiosis is associated with various inflammatory diseases such as IBD, RA, and osteoporosis [11]. The World Health Organisation (WHO) recommendations state that probiotics are live bacteria that, when taken in sufficient quantities, have positive health effects [12]. Several studies, including ours, have demonstrated the beneficial effects of probiotics in osteoporosis [13,14]. However, their use is limited in immunocompromised individuals, making postbiotics a promising alternative. Postbiotics are small metabolites produced by the gut microbiota (GM) as metabolic by-products of the undigested food components, such as short-chain fatty acids (SCFAs), indole derivatives, and tryptophan metabolite (indole acetic acid). The GM not only metabolises exogenous dietary components but also acts on the primary bile acids (BAs) such as cholic acid and chenodeoxycholic acid, which are synthesised from cholesterol in the liver and transported to the gut, where most of them are resorbed and the remaining are converted to the secondary BAs by the 7α-dehydroxylation activity of GM. The secondary BAs, deoxycholic acid (DCA) and lithocholic acid (LCA), are further modified by the hydroxysteroid dehydrogenase (HSDH) activity of the GM [15]. The dysbiosis leads to decreased levels of secondary BAs in the gut [16]. BAs function through certain receptors, such as G-protein-coupled bile acid receptor 5 (TGR5) and farnesoid X receptor (FXR). It has been reported that secondary BAs promote Treg differentiation while inhibiting the Th17 cells in the spleen and intestine, alleviating intestinal inflammation [17]. Additionally, BAs stimulate intestinal stem cell receptors to repair the intestinal epithelium, promoting the differentiation and regeneration of epithelial cells [18]. In autoimmune uveitis, a recent study found that exogenous DCA suppressed Th1, Th17, and dendritic cells and decreased the severity of the disease [19]. BAs thus play a crucial role in immune regulation and maintaining homeostasis, and therefore, modulating BA activity may offer a promising strategy to improve osteoporosis. However, the role of DCA in mitigating inflammatory bone loss has not been elucidated until now. In this study, we demonstrate that DCA alleviates LPS-induced inflammatory bone loss by regulating bone remodelling and modulating the gut–bone axis. Collectively, our findings highlight the protective effects of DCA on osteoblast and osteoclast function, gut barrier integrity, and overall bone health.

### 2.0 Materials & Methods

### 2.1 Reagents and antibodies

Sigma (USA) provided the Acid Phosphatase Leukocyte (TRAP) kit, while Cayman Chemical supplied the deoxycholic acid (DCA, CAS 83-44-3). ProSpec provided the recombinant murine M-CSF (CYT-308) and RANKL (CYT-334). Gibco (USA) supplied culture media, such as RPMI-1640 and α-MEM, whereas Falcon provided basic tissue culture plasticware. Promega (USA) provided the RT-PCR master mix. The cDNA synthesis kit, gene-specific primers, Alizarin Red S (A5533, Sigma, USA), and the ALP staining kit (NBT/BCIP, 34042, Thermo, USA) were among the staining and differentiating tools.

Antibodies were obtained from multiple suppliers: GAPDH (2118S), Cathepsin K (57056S), c-Fos (2250S), NFATc1 (4389S), Runx2 (12556S), and FXR (72105S) from Cell Signaling Technology (USA); HRP-conjugated anti-mouse (58802S) and anti-rabbit (7074) secondary antibodies from Cell Signaling Technology (USA); and the TGR5 antibody (PA5-23182) from Thermo Fisher Scientific (Invitrogen, USA). Additional reagents included a phosphatase inhibitor (2850, Sigma, USA), RIPA buffer (89900, Thermo, USA), a protease inhibitor cocktail (5871S, Cell Signaling Technology), and the BCA protein quantification kit (A55860, Thermo, USA).

### 2.2 LPS-induced inflammatory bone loss

C57BL/6 mice aged 8–10 weeks were used in for in vitro and in vivo studies. The animals were kept in the animal facility of the All India Institute of Medical Sciences (AIIMS), located in New Delhi, India, under specified pathogen-free (SPF) conditions. Mice were randomized to one of three groups for the in vivo experiments: Control (healthy), LPS-induced, and LPS + DCA. An intraperitoneal dose of 5 mg/kg of LPS was given on days 0 and 4. At the end of the experimental period (11 days), mice from all groups were euthanised, and blood, bones, and lymphoid organs were collected for osteoimmune parameter analysis. All procedure were carried out with the Institutional Animal Ethics Committee of AIIMS, New Delhi,’s clearance (Protocol No. 338/IAEC-1/2021).

### 2.3 Quantitative PCR (q-PCR)

RNA was isolated from the bone marrow cells using the RNeasy Mini Kit (Qiagen, USA) in accordance with the manufacturer’s instructions. A Bio Analyser (Agilent Technologies, Singapore) was used to determine the quality of the RNA, and samples with an RNA Integration Number (RIN) value more than seven underwent further processing. Additionally, a spectrophotometer (Nanodrop Technologies, USA) was used to measure the amount of RNA. A Thermo Scientific cDNA synthesis kit was utilized for cDNA conversion. Custom primers for c-Fos, Nfatc1, TRAP, RANKL, osteocalcin, Runx2, OPG, TGR5, FXR, claudin-1, and occludin were used to amp up triplicate quantities of cDNA from each group. The arithmetic mean of the GAPDH housekeeping gene was used to normalize the results. For each reaction, 25 ng of cDNA was used in each well that included the primers and 2X SYBR green PCR master mix (Promega, USA). Last but not least, threshold cycle data were converted to relative gene expression after normalization.

### 2.4 Micro-computed tomography (µ-CT) measurements

Following animal dissection, the hind limbs were gathered, and all of the muscles were cleared of any bones. The storage solution for these bone samples was 4% paraformaldehyde. As previously mentioned, µ-CT scanning and analysis were carried out utilising an in vivo X-ray SkyScan 1076 scanner (Aartselaar, Belgium). In summary, the samples were oriented correctly in the sample holder and scanned at 50 kV, 204 mA, using a 0.5-mm aluminum filter. The NRecon program was utilised for the rebuilding procedure. 100 slices of ROI were removed from the secondary spongiosa after rebuilding, spaced 1.5 mm from the distal end of the development plate. Further, CTAn software was used to process ROI and compute the micro-architectural parameters of the bone samples. Trabecular thickness (Tb.Th), trabecular number (Tb. N), endosteal perimeter (E.Pm), trabecular separation (Tb.Sp), bone volume/tissue volume (BV/TV), and other 3D-histomorphometric values were acquired. The BMD of the femur and tibia were calculated using the volume of interest of µ-CT images performed for trabecular and cortical regions. As a calibrator, hydroxyapatite phantom rods with a diameter of 4 mm and known BMDs of 0.25 and 0.75 g/cm³ were used to test BMD.

### 2.5 Osteoclast Differentiation and TRAP Staining

Bone marrow cells (BMCs) were harvested from the femurs and tibias of 8–12-week-old C57BL/6J mice by flushing with α-MEM supplemented with 10% heat-inactivated FBS. Following red blood cell lysis using 1× RBC lysis buffer, the cells were cultured overnight in T-25 flasks containing endotoxin-free α-MEM with 10% heat-inactivated FBS and M-CSF (35 ng/mL). The next day, non-adherent cells were collected, and 5 × 10^4 cells were seeded into 96-well plates in α-MEM containing M-CSF (30 ng/mL) and RANKL (100 ng/mL), in the presence or absence of DCA, and maintained for four days. On day 3, half of the culture medium was replaced with fresh α-MEM supplemented with the respective factors.

At the end of culture, osteoclast differentiation was assessed using tartrate-resistant acid phosphatase (TRAP) staining according to the manufacturer’s protocol. Briefly, cells were rinsed three times with PBS, fixed for 10 min at room temperature in a solution of citrate, acetone, and 3.7% formaldehyde, and then washed twice with PBS. TRAP staining was performed by incubating the cells with TRAP substrate solution at 37 °C in the dark for 5–15 min. Multinucleated TRAP-positive cells containing ≥3 nuclei were identified as osteoclasts, which were imaged using inverted microscopes (Eclipse TS100, Nikon; EVOS, Thermo Fisher Scientific). Osteoclast surface area was quantified with ImageJ software (NIH, USA).

### 2.6 Alkaline Phosphatase (ALP) & Alizarin Red S staining

For alkaline phosphatase (ALP) staining, cells were rinsed with PBS and fixed in 4% paraformaldehyde for 30 min at room temperature. After fixation, the paraformaldehyde was discarded, and cells were incubated with BCIP/NBT substrate solution at 37 °C in the dark for 15 min. The reaction was stopped by washing with PBS, and stained cultures were visualized and imaged using an EVOS XL Core microscope (Thermo Fisher Scientific, USA).

For alizarin red staining, culture medium was aspirated and cells were washed twice with PBS. Cells were then fixed with 10% formaldehyde (100 µL per well) for 30 min at room temperature, followed by two additional PBS washes. A 2% alizarin red solution prepared in distilled water (50 µL per well) was added, and plates were incubated at 37 °C until calcium deposits were visibly stained. To quantify mineralization, the bound dye was eluted with 100 µL of a solution containing 10% acetic acid, 20% methanol, and 70% distilled water, followed by incubation for 15 min. Absorbance was recorded at 450 nm using a BioTek Synergy H1 spectrophotometer/fluorimeter.

### 2.7 F-Actin Ring Formation Assay

Osteoclast cytoskeletal organization was evaluated using phalloidin staining. Bone marrow– derived osteoclast precursors were cultured in 96-well plates, and on day 4 cells were processed for F-actin visualization. Cultures were rinsed twice with PBS, fixed in 4% paraformaldehyde for 20 min, and permeabilized with 0.1% Triton X-100 for 5 min. To reduce nonspecific staining, cells were incubated with 1% BSA for 30 min before labeling with FITC-conjugated phalloidin for 30 min at room temperature in the dark. Nuclear counterstaining was performed with DAPI (1 µg/mL, 5 min, dark). F-actin ring structures were examined under a fluorescence microscope (Eclipse Ti, Nikon).

### 2.8 Enzyme-Linked Immunosorbent Assay

Serum levels of IL-2, IL-4, TGF-β, IL-10, IL-17, TNF-α, IFN-γ, IL-6, and RANKL were measured using commercially available ELISA kits, following the manufacturers’ protocols.

### 2.9 Immunofluorescence Staining

Osteoclasts cultured in 96-well plates were processed for immunofluorescence on day 4. Cells were fixed with 4% paraformaldehyde for 20–30 min at room temperature, rinsed with PBS, and permeabilized with 0.2% Triton X-100 for 2 h at 37 °C. Following three washes in PBS containing 0.05% Triton X-100, nonspecific binding was blocked using 2% goat serum in wash buffer for 1 h at room temperature. Cultures were then incubated overnight at 4 °C with primary antibodies specific to the proteins of interest. After three PBS washes, fluorophore-conjugated secondary antibodies were applied, and nuclei were counterstained with DAPI.

### 2.10 Histological evaluation of osteolysis

Femurs were fixed in 10% formalin at room temperature, followed by removal of surrounding soft tissue. Bones were decalcified for 3 weeks in a commercial decalcifying solution (Sigma, D0818), dehydrated, and embedded in paraffin. Serial sections (5 μm) were prepared, stained with TRAP and fast green at 37 °C according to the manufacturer’s instructions, and examined under a light microscope (Euromax) at 40× magnification. TRAP-positive osteoclasts were quantified in six randomly selected fields. Intestinal tissues were processed in parallel, sectioned, and stained with hematoxylin and eosin (H&E) for histological evaluation.

### 2.11 Western Blotting

Osteoclasts derived from mouse BMSCs and cultured with or without DCA were lysed in RIPA buffer containing protease inhibitors. Protein concentrations were quantified using the BCA method. Equal amounts of protein were resolved on 12% SDS–PAGE gels and transferred onto nitrocellulose membranes. Membranes were blocked with 5% BSA and incubated overnight at 4 °C with primary antibodies specific to the target proteins. Following PBS washes, membranes were probed with HRP-conjugated secondary antibodies for 2 h at room temperature. Protein bands were visualized using an enhanced chemiluminescence detection system, and band intensities were quantified using ImageJ software. GAPDH served as the loading control.

### 2.12 Statistical analysis

Data are presented as mean ± SEM (n = 6). Group comparisons were performed using one-way ANOVA, followed by Student’s t-test (paired or unpaired, as appropriate). Statistical significance was defined as *p* ≤ 0.05, with thresholds indicated as p < 0.05 (), p < 0.01 (), and p < 0.001 ().

## 3. Results

### 3.1 DCA promotes Osteoblastogenesis of the BMCs

In order to ascertain DCA’s osteogenic capacity in ex vivo settings, BMCs were cultivated in osteogenic media with or without DCA at different doses (1 μM and 5 μM) **(Figure 1A)**. On 14^th^ day, cells were fixed and processed for ALP staining. Interestingly, DCA markedly increased osteoblastogenesis in a concentration-dependent way. **(Figure 2A).** Further, the effect of DCA on bone mineralisation was evaluated. To accomplish the same, BMCs were cultured with DCA in the presence of osteogenic differentiation media. In order to stain the calcium nodules, cells were prepared for alizarin red staining on the twenty-first day. The results show that DCA considerably increased bone mineralization. (**Figure 2B),** which is further confirmed by the quantification of the alizarin red staining, which showed a marked increase in mineral deposition upon DCA treatment **(Figure 2B-C).** Altogether, our results indicated that DCA significantly promotes the osteoblastogenesis of the BMCs.

**Figure 1.**
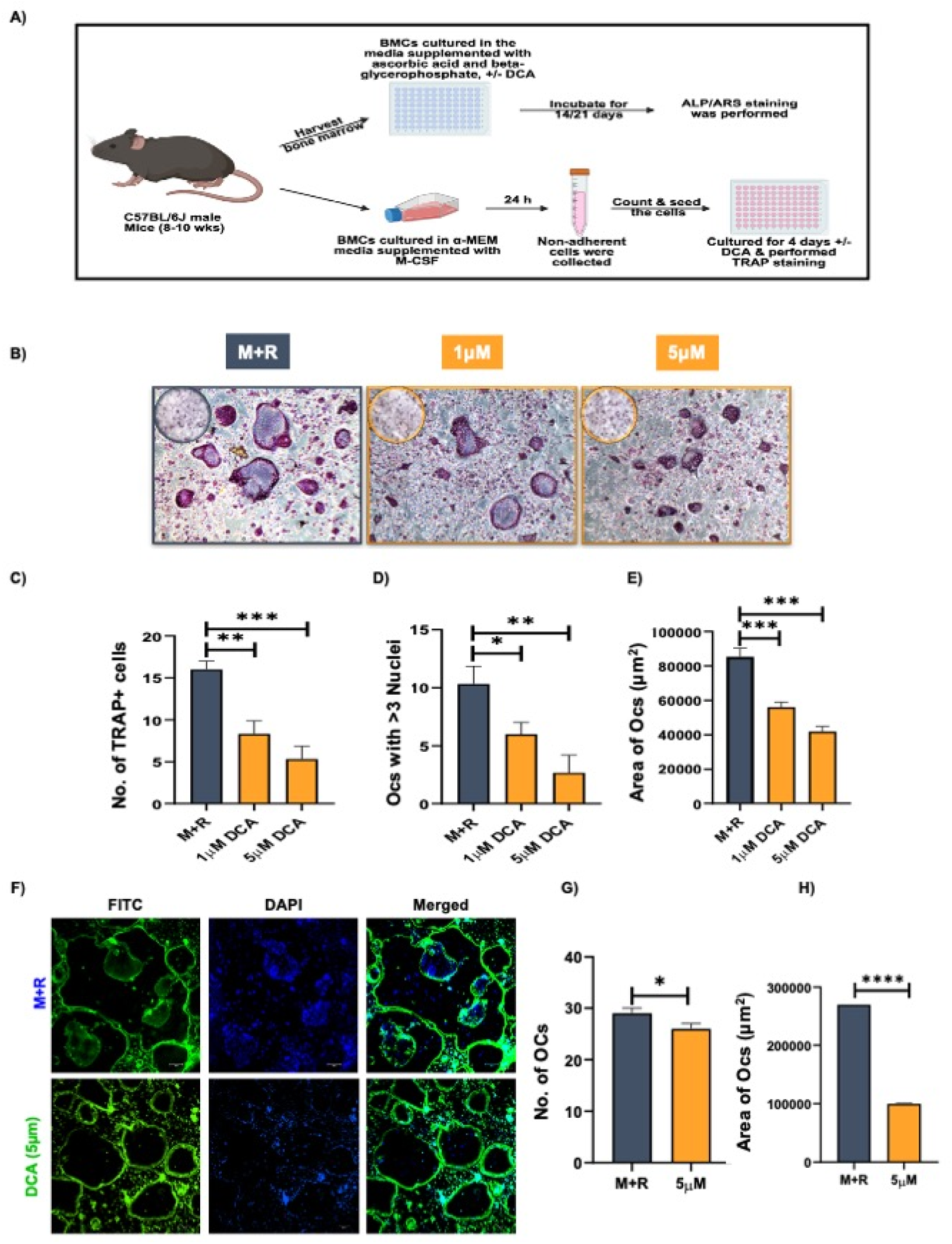
DCA treatment suppresses osteoclastogenesis. **A)** showing methodology of osteoclast and osteoblast. **B)** DCA significantly inhibited the generation of multinucleated osteoclasts. **C)** Bar graph representing the number of TRAP-positive cells. **D)** number of TRAP-positive cells with more than 3 nuclei. **E)** area of multinucleated TRAP-positive cells. **F)** F-actin and nuclei stained with phalloidin and DAPI, respectively. **G)** Bar graph representing the number of multinucleated cells. **H)** area of acting rings.

**Figure 2.**
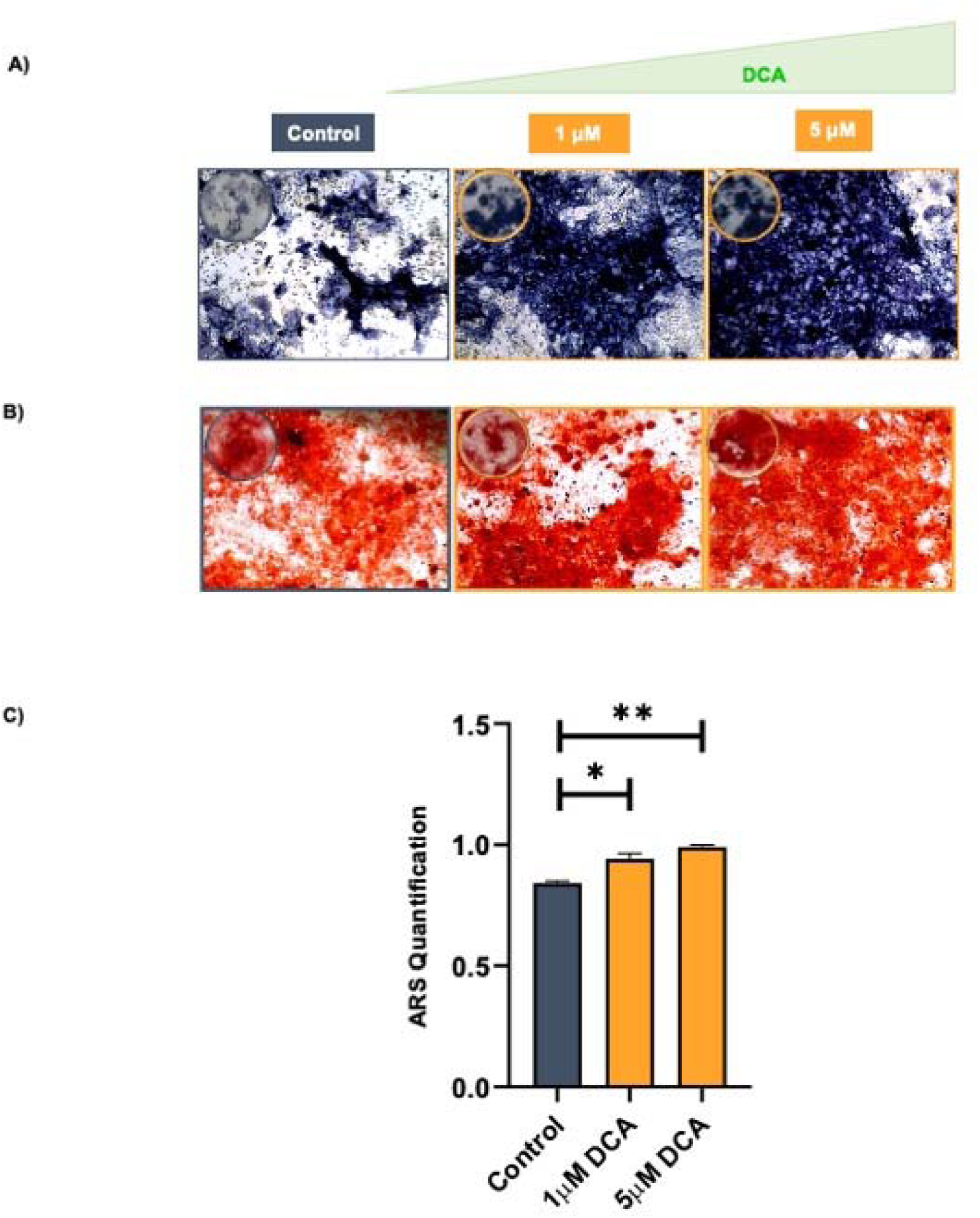
DCA treatment enhanced osteoblastogenesis and mineralisation. **A)** BMCs were cultured in the media supplemented with ascorbic acid (50μg/ml) and beta-glycerophosphate (10mM) with or without DCA, for 14 or 21 days. **B)** DCA treatment enhanced osteoblastogenesis in a dose-dependent manner. **C)** DCA enhanced mineralization in BMCs in a dose-dependent manner. **D)** Graphical representation showing the quantification of ARS.

### 3.2 DCA suppressed RANKL-induced Osteoclast differentiation

Next, we assessed how DCA affected the differentiation of osteoclasts. This was accomplished by cultivating BMCs in the whole α-MEM medium supplemented with MCSF (30 ng/ml) and RANKL (100 ng/ml) with or without varying DCA doses (1 and 5 µM). Interestingly, the number of TRAP-positive cells and cells with more than three nuclei were reduced, indicating that DCA effectively suppressed osteoclastogenesis with efficacy increasing according to concentration **(Figure 1C-D).** Additionally, the area of osteoclasts in the DCA groups was significantly smaller than in the LPS-treated group. **(Figure 1E).** Furthermore, DCA administration dramatically reduced the area and number of F-actin rings, suggesting that DCA suppresses both osteoclast development and osteoclast functional activity **(Figure 1F-G).**

### 3.3 DCA modulates Osteoclast and Osteoblast-specific genes and proteins

Our previous results demonstrated that DCA promotes *in vitro* bone formation while inhibiting osteoclast function. To further investigate its molecular effects, we performed gene and protein expression analyses **(Figure 3A).** DCA treatment significantly enhanced the mRNA expression of FXR and TGR5 genes, the BA receptors **(Figure 3B)**. DCA treatment suppressed the expression of key osteoclastogenic genes, including c-Fos, CTSK, NFATc1, and TRAP **(Figure 3C).** It consistently inhibited the protein levels of c-Fos, CTSK, and NFATc1 during osteoclast differentiation **(Figure 3E).** Immunofluorescence analysis further confirmed that DCA reduced osteoclastogenic gene expression **(Figure 3F)**. Conversely, DCA treatment significantly upregulated the osteogenic marker Runx2 at protein levels in BMCs **(Figure 3D).** Collectively, these findings indicate that DCA attenuates osteoclast development and function by downregulating osteoclast-specific genes and proteins.

**Figure 3.**
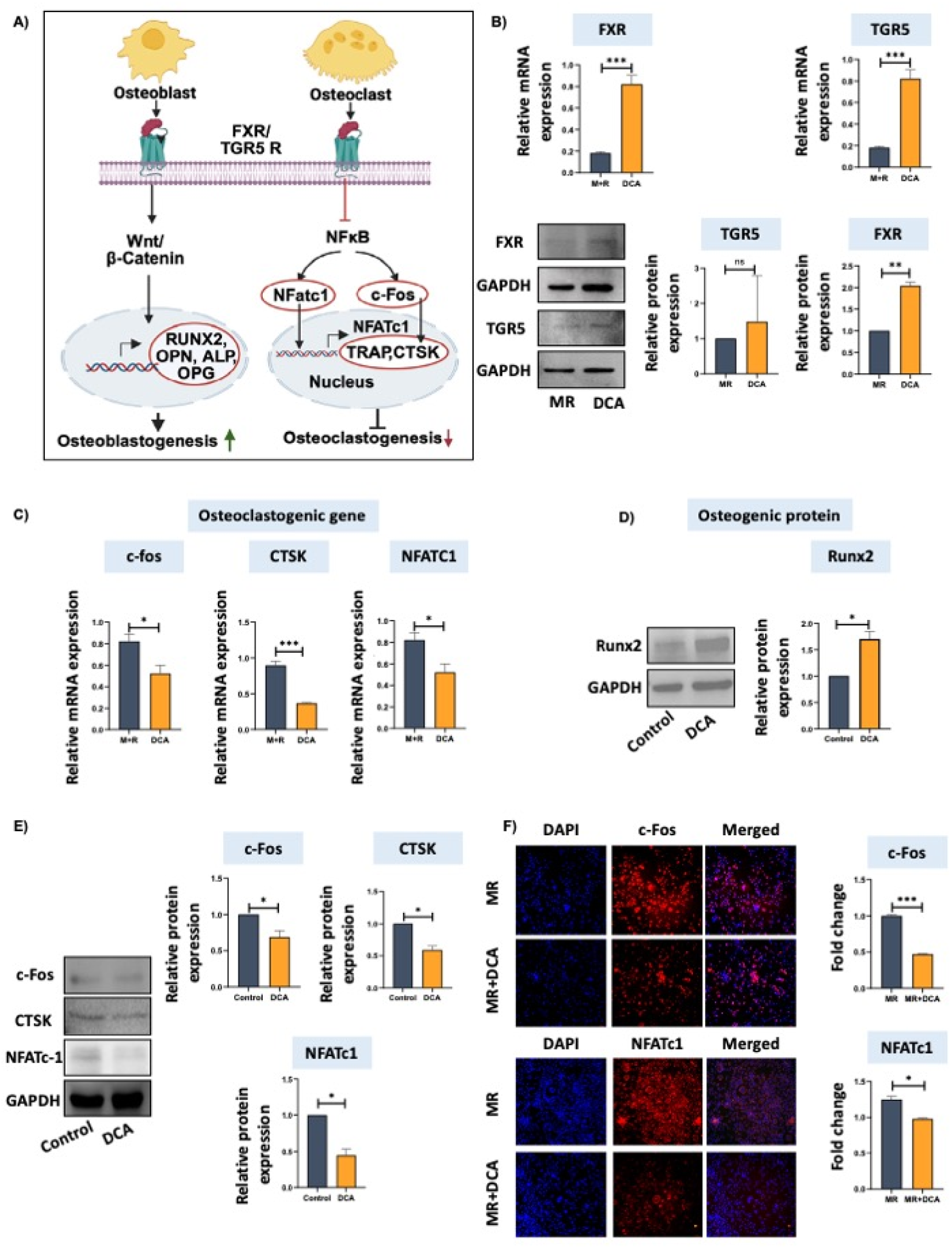
DCA modulates bone remodeling via FXR and TGR5 signaling. **Fig A)** Showing osteoblast and osteoclast signalling. **B)** relative mRNA and protein expression of FXR and TGR5, normalized with GAPDH. **C)** enhanced Runx2 expression after DCA treatment. **D)** The relative intensity mrNA expression of osteoclastogenic genes c-for, CTSK and NFATC1 was reduced by DCA treatment. **E)** DCA treatment inhibited expression of osteoclastogenic proteins, C-Fos, NFATc1, normalised with GAPDH. **F)** Immunofluorescence staining demonstrated decreased expression of osteoclastogenic genes.

Previous studies have shown that DCA mediates its effects through the bile acid receptors FXR and TGR5 [20]. Consistent with this, we observed that DCA treatment increased both mRNA and protein expression of FXR and TGR5 **(Figure 3B).** Taken together, these results suggest that DCA exerts osteoprotective effects by simultaneously promoting osteoblastogenesis and inhibiting osteoclastogenesis through FXR and TGR5 signalling pathways.

### 3.4 DCA attenuates LPS-induced Inflammatory Bone Loss *in vivo*

To investigate the effects of DCA on inflammatory bone loss *in vivo*, we employed an LPS-induced osteolysis model in C57BL/6J mice, in which LPS was administered intraperitoneally. Micro-CT analysis showed that LPS treatment markedly impaired the trabecular bone microarchitecture of the femur and tibia compared with control mice **(Figure 4A).** Notably, DCA treatment reversed LPS-induced bone loss, as evidenced by significant improvements in BMD, bone volume fraction (BV/TV), and trabecular number (Tb. N), along with reduced trabecular separation (Tb. Sp) **(Figure 4B–E).**

**Figure 4.**
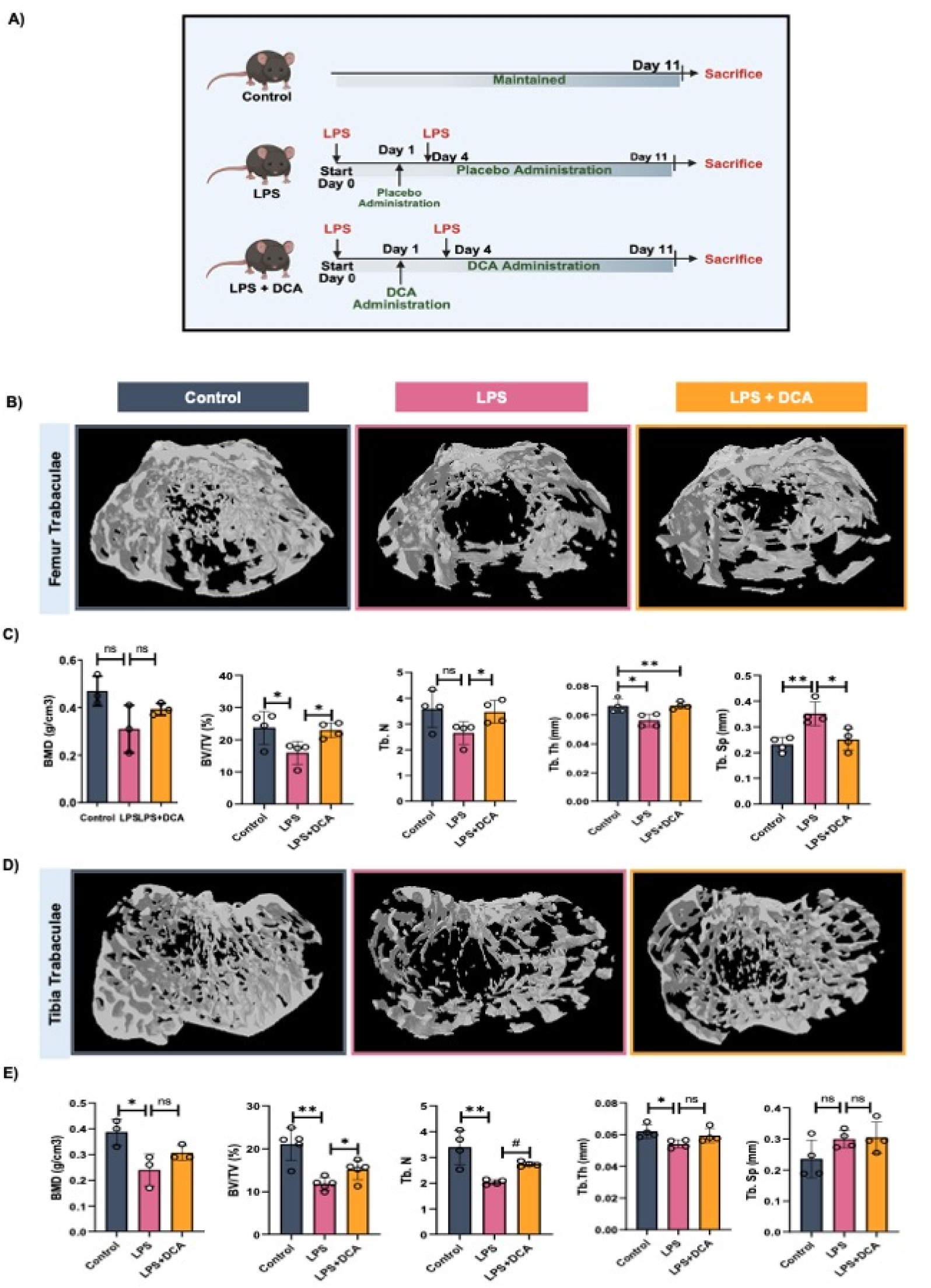
DCA administration improves trabecular bone microarchitecture. 3D uCT reconstruction of femur trabecular, and tibia trabecular of all groups. **A)** Development of LPS-induced inflammatory bone loss model. **B)** Bone micro-architecture of femur trabecular. **C)** Histomorphometric parameters of femur trabecular. **D)** Bone micro-architecture of tibia trabecular. **E)** Histomorphometric parameters of tibia trabecular. BV/TV, bone volume/tissue volume ratio; Tb. Th., trabecular thickness; Tb. Sp., trabecular separation; Tb.N., trabecular number. The results were evaluated by ANOVA with subsequent comparisons by Student’s t-test for paired or non-paired data. Values are reported as mean ± SEM. The above graphical representations are indicative of one independent experiment, and similar results were obtained in two different independent experiments with n = 6. Statistical significance was considered as p ≤ 0.05 with respect to indicated mouse groups.

In addition, DCA treatment significantly restored cortical bone parameters altered by LPS, including total cross-sectional perimeter (T.Pm), cross-sectional thickness (Cs.Th), endosteal perimeter (E.Pm), and medullary area (M.Ar) in both femur and tibia **(Figure 5A–D).** These results indicate that DCA effectively ameliorates LPS-induced deterioration of trabecular and cortical bone.

**Figure 5.**
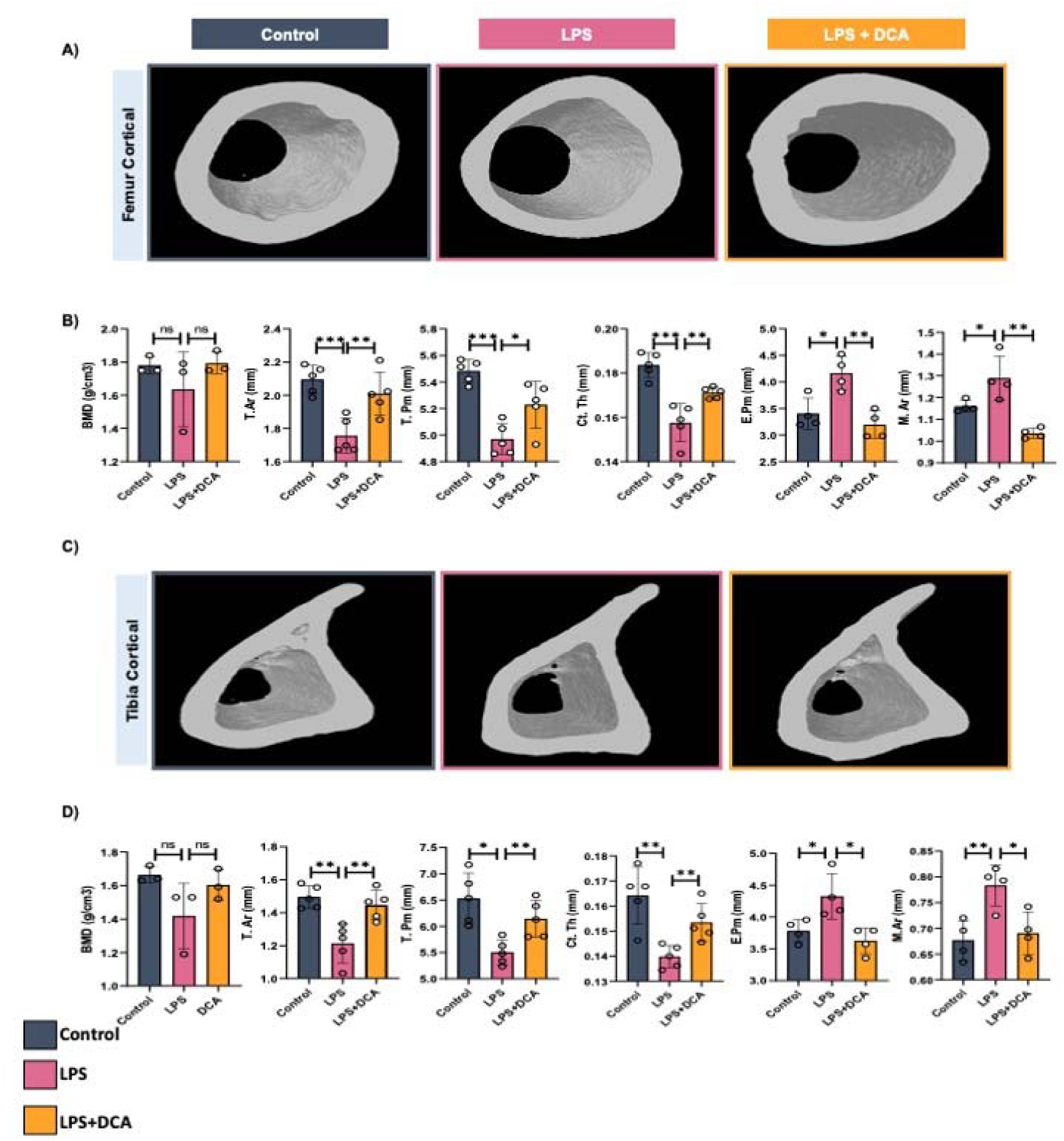
DCA administration improves cortical bone microarchitecture. **3D** μ**-CT reconstruction of femoral cortical and tibia cortical of all groups. A)** Bone micro-architecture of femur cortical. **B)** Histomorphometric parameters of femur cortical. **C)** Bone micro-architecture of tibia cortical. **E)** Histomorphometric parameters of tibia cortical. Histomorphometric parameters of tibia cortical. **D)** Histomorphometric parameters of tibia cortical. Tt. Ar., total cross-sectional area; T. Pm., total cross-sectional perimeter; Ct. Th., cortical thickness. The results were evaluated by ANOVA with subsequent comparisons by Student’s t-test for paired or non-paired data. Values are reported as mean ± SEM. The above graphical representations are indicative of one independent experiment, and similar results were obtained in two different independent experiments with n = 6. Statistical significance was considered as p ≤ 0.05 (*p ≤ 0.05, **p ≤ 0.01) with respect to indicated mouse groups.

### 3.5 DCA suppressed Osteoclastogenesis in LPS-treated mice

To further assess the effect of DCA on osteoclastogenesis, bone marrow cells isolated from control, LPS-treated, and LPS + DCA-treated mice were cultured in osteoclastogenic media supplemented with M-CSF and RANKL. TRAP staining performed on day 4 revealed a significant reduction in the number of TRAP-positive cells and cells with >3 nuclei in the DCA-treated group compared with LPS-treated mice **(Figure 6A–C).** In addition, DCA treatment markedly decreased the area of multinucleated cells **(Figure 6D).** Further, we evaluated the effect of DCA on the expression of osteoclastogenic and anti-osteoclastogenic genes in bone marrow cells. DCA treatment significantly suppressed osteoclastogenic markers, including c-Fos, NFATc1, RANKL, TRAP, and the RANKL/OPG ratio **(Figure 6E).** Conversely, DCA markedly upregulated osteogenic gene expression **(Figure 6F).** These results suggest that DCA not only inhibits osteoclast differentiation but also promotes osteoblast activity, thereby restoring bone remodelling under inflammatory conditions.

**Figure 6.**
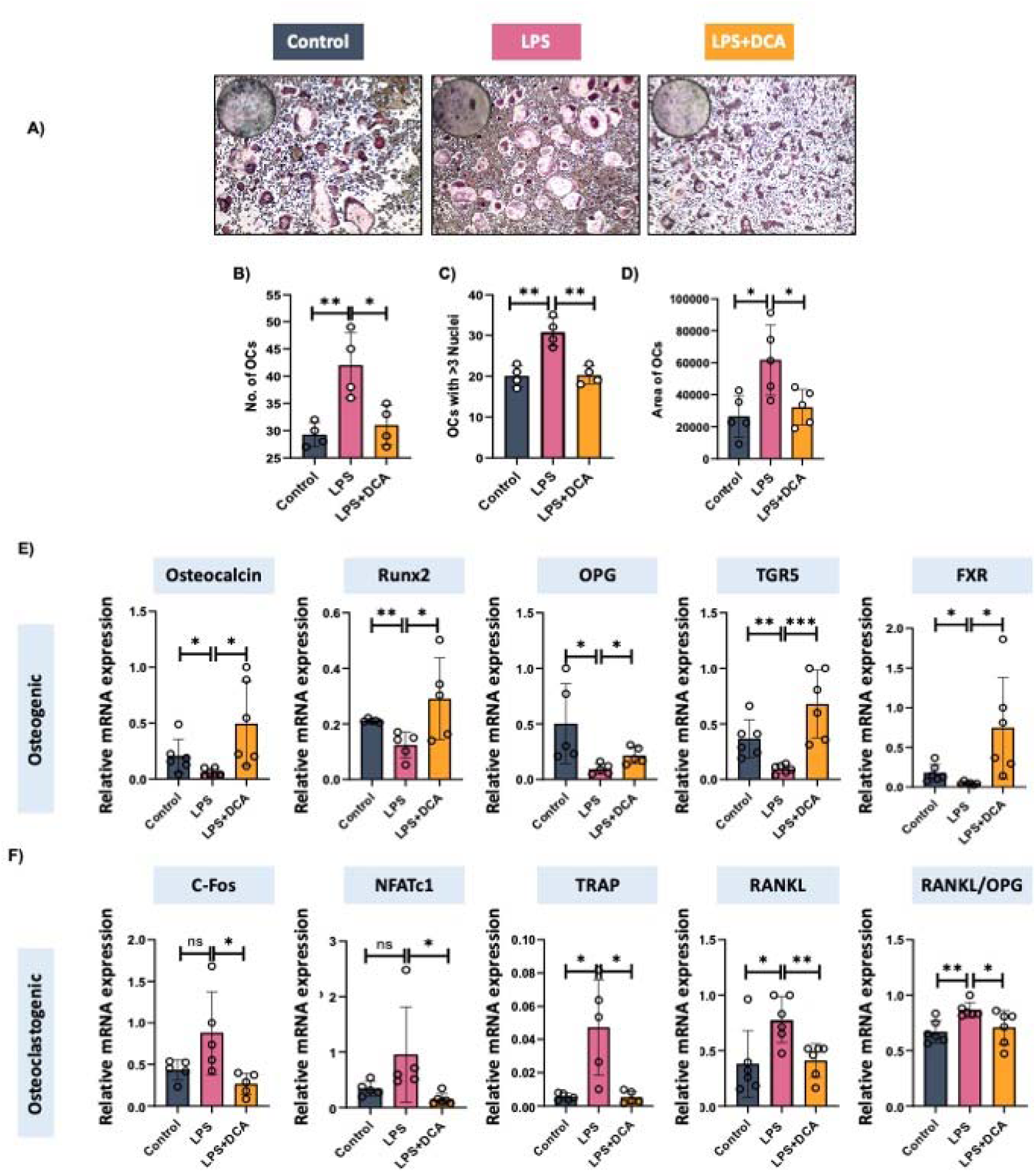
DCA administration prevents osteoclastogenesis. **A)** Osteoclast culture of control, LPS and DCA-treated mice. **B)** Bar graph representing TRAP-positive cells. **C)** number of multinucleated cells. **D)** area of osteoclast. **E)** expression of osteoclastogenic genes, including c-Fos, NFATc1, TRAP, RANKL and RANKL/OPG ratio. **F)** expression of anti-osteoclastogenic genes, osteocalcin, Runx2, and OPG.

### 3.6 DCA augments Bone health via modulating cytokine secretion

Inflammation promotes osteoclastogenesis by enhancing the secretion of pro-inflammatory cytokines while suppressing anti-inflammatory cytokines [21]. To evaluate whether DCA influences this process, we analysed the cytokine profile and observed that DCA treatment effectively reversed the alterations induced by LPS **(Figure 7A–B).** Collectively, these findings suggest that DCA modulates immune cell function in LPS-treated mice, restoring the balance between pro- and anti-inflammatory cytokines, thereby reducing osteoclastogenesis.

**Figure 7.**
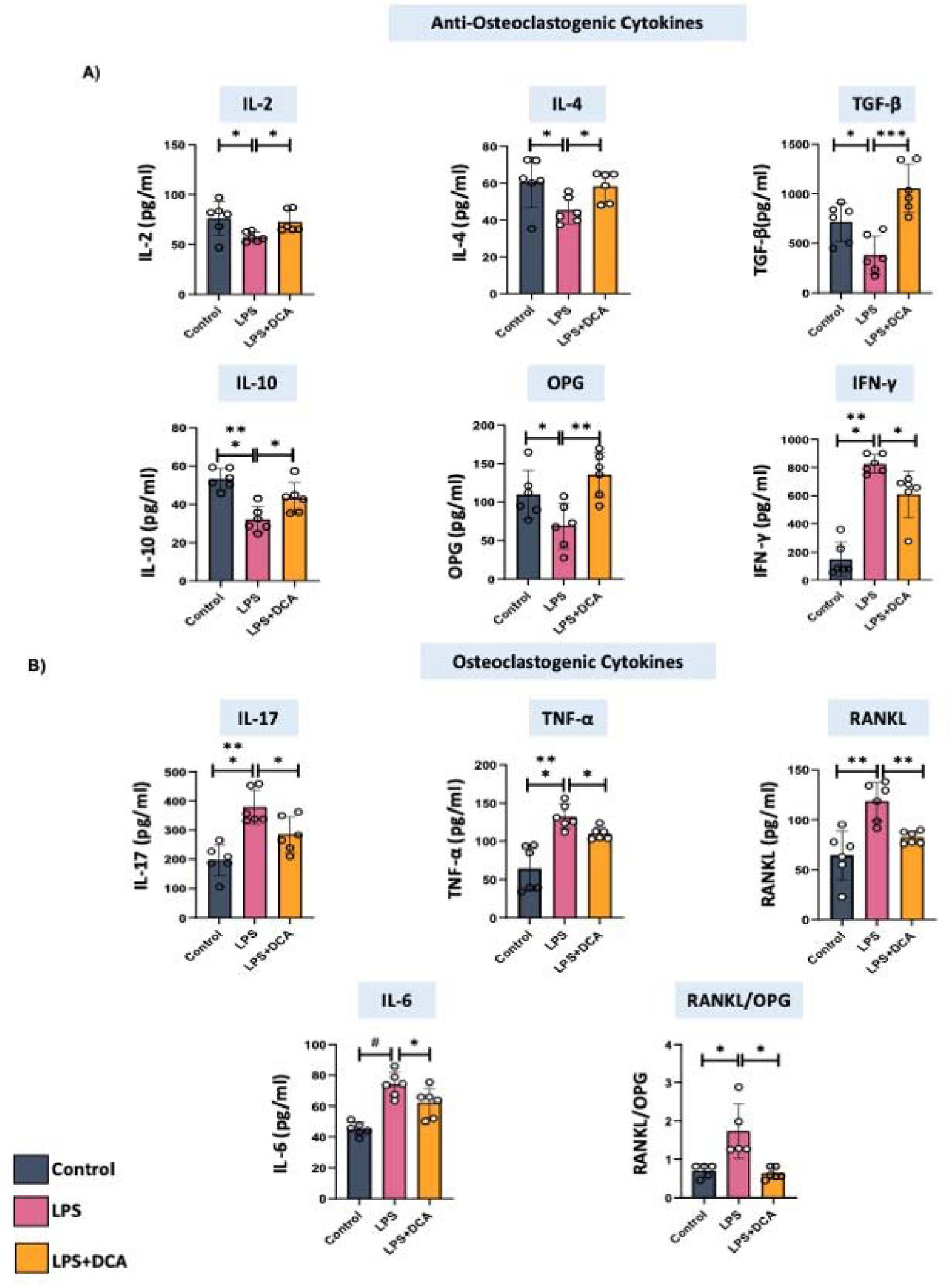
DCA Attenuates LPS-Induced Cytokine Imbalance. **A)** Anti**-**Osteoclastogenic, **B)** Osteoclastogenic cytokines were analysed in serum samples of mice. Data were analysed using one-way ANOVA followed by Student’s *t*-test for paired or unpaired comparisons, as appropriate. Results are presented as mean ± SEM (n = 6), and similar outcomes were observed in two independent experiments. Statistical significance was defined as *p* ≤ 0.05. Significance levels are indicated as follows: *p* ≤ 0.05 (*), p < 0.01 (), and p* ≤ *0.001 (),* relative to the control group.

### 3.7 DCA promotes gut health

Prior research conducted by our team and others has demonstrated a robust correlation between bone homeostasis and gastrointestinal health.[22]. Thus, we next examined the role of DCA in maintaining bone homeostasis via regulating gut health. Histological analysis revealed that LPS treatment caused marked intestinal damage, whereas DCA administration restored normal gut architecture **(Figure 8A).** Consistent with this, LPS treatment was associated with increased gut permeability, as evidenced by elevated expression of IL-23 and reduced expression of tight junction proteins such as claudin and occludin **(Figure 8B).** Importantly, DCA treatment reversed these alterations, thereby preventing leaky gut. Since gut permeability is closely linked to dysbiosis, we further performed gut microbiome analysis in all the groups of mice. We found a reduction in the relative abundance of Bacteroidota and Patescibacteria and enhanced Firmicutes abundance in the LPS-treated mice group, which was restored by DCA treatment in mice **(Figure 8D-F)**.

**Figure 8.**
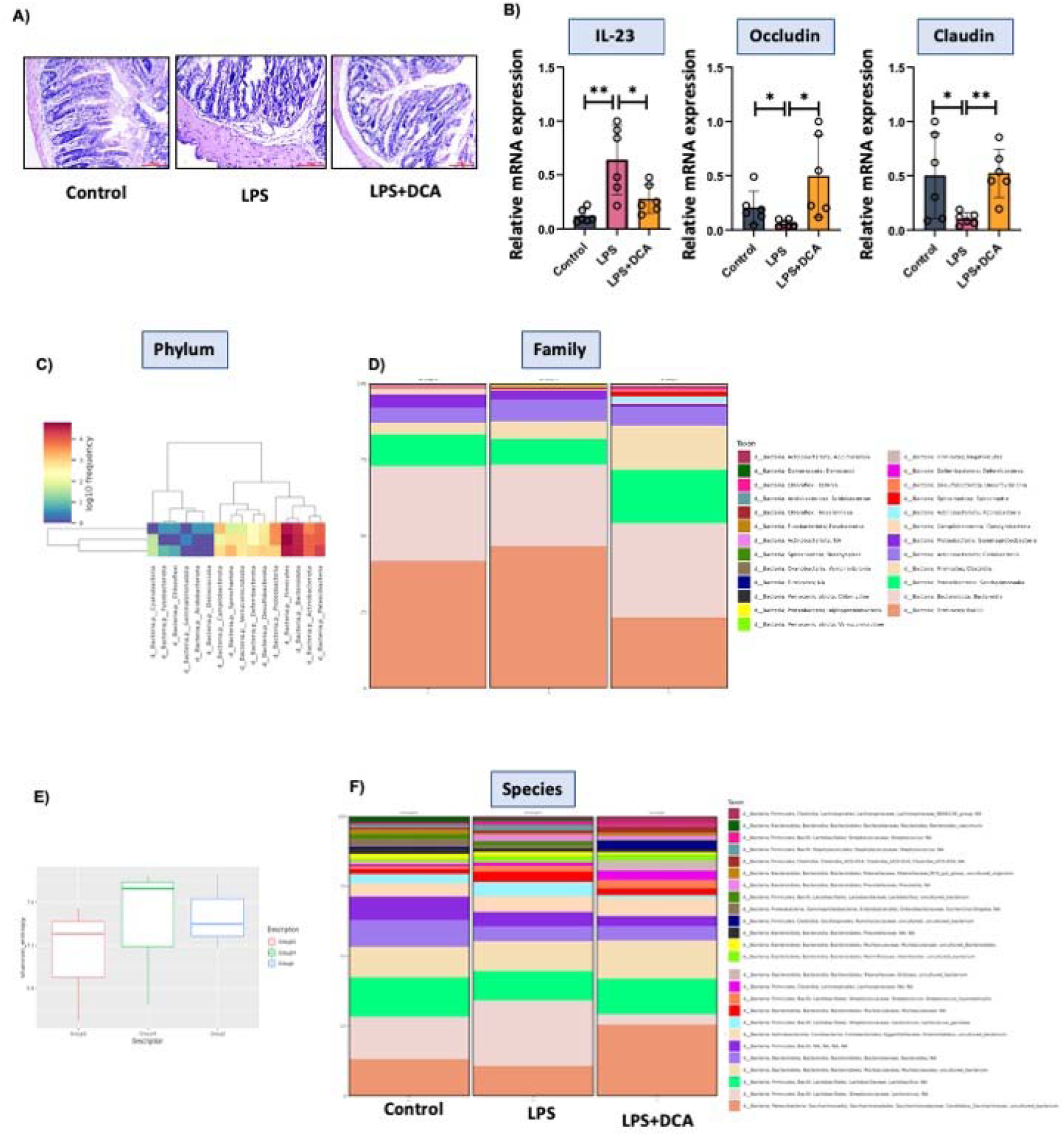
DCA promotes gut intigrity and microbial balance. **A)** H&E of small intestine tissues, **B)** tight junction proteins, occluding, claudin and IL-23**. C)** heatmap, **D)** Bar graph showing family and genus. **E)** Shannon diversity index.

## 4. Discussion

Osteoblasts and osteoclasts work together in a tightly controlled process called bone remodelling to preserve mineral balance and skeletal integrity[23,24]. Disruption of the homeostatic balance between osteoclasts and osteoblasts contributes to diseases such as osteoporosis, highlighting the importance of therapeutically targeting bone remodelling to preserve bone health. In the present study, we identified DCA as a modulator of bone remodelling with dual pharmacological properties. DCA markedly promotes the osteoblastogenesis and mineralisation of the BMCs. DCA further suppressed the RANKL-induced osteoclast development. Quantitative analysis showed that DCA significantly decreased the quantity, size, and multinucleation of osteoclasts. Additionally, functional experiments showed that DCA inhibited bone resorption activity and F-actin ring formation, indicating decreased osteoclast functional activity. Furthermore, the expression of important osteoclastogenic transcription factors was downregulated by DCA at the molecular level. Western blotting and qPCR verified that DCA treatment markedly reduced RANKL-induced transcriptional activity and c-Fos and NFATc1 protein expression. These findings offer a molecular explanation for the observed inhibition of osteoclastogenesis, given the critical roles that c-Fos and NFATc1 play in promoting osteoclast lineage commitment [25,26]. In a mouse model, fed with DCA successfully prevented LPS-induced bone loss *in vivo*. In comparison to controls, DCA-treated mice’s femoral and tibial areas showed significant improvements in both trabecular and cortical bone microarchitecture, according to micro-computed tomography (μ-CT) studies. Importantly, DCA prevented bone resorption while simultaneously encouraging osteoblast development, which increased bone mass overall. Taken together, these findings establish DCA as a potential therapeutic candidate for bone loss disorders. By simultaneously stimulating osteoblastogenesis and inhibiting osteoclastogenesis, DCA restores bone remodelling balance and preserves skeletal microarchitecture under pathological conditions.

BAs exert diverse physiological effects not only as detergents facilitating lipid absorption but also as signalling molecules that regulate inflammation, metabolism, and tissue homeostasis. The activation of bile acid receptors, including nuclear receptors and G protein-coupled receptors (GPCRs), is primarily responsible for these immunomodulatory effects [27]. The two main receptors are the membrane-bound GPCR TGR5 and the ligand-activated nuclear receptor FXR. Both receptors have been linked to controlling osteoimmune interactions and inhibiting inflammatory signalling pathways. Prior research has shown that in preclinical models of vascular inflammation and experimental autoimmune encephalomyelitis (EAE), pharmacological stimulation of FXR and TGR5 reduces inflammatory responses [28,29]. Importantly, it has been demonstrated that FXR and TGR5 activation restricts osteoclast development and prevents bone loss in estrogen-deficient conditions [30]. These results demonstrate the critical role of bile acid signalling in maintaining bone homeostasis.. Secondary bile acids, such as DCA and LCA, preferentially bind and activate TGR5, while primary bile acids, such as CDCA and CA, are the strongest endogenous activators of FXR [31]. Our investigation showed that oral treatment of DCA dramatically increased the expression levels of both FXR and TGR5 in bone tissue, which is consistent with this receptor-ligand paradigm. This suggests that DCA may exert its dual anti-inflammatory and anti-resorptive effects by simultaneously activating nuclear receptor–mediated transcriptional pathways and GPCR-dependent signalling cascades. All of these findings point to receptor-mediated signalling via FXR and TGR5 as a major mechanistic pathway that underlies DCA’s osteo-protective effects. Through the upregulation of these BA receptors, DCA preserves bone microarchitecture in pathological situations by inhibiting osteoclastogenesis and dampening pro-inflammatory signals.

It is becoming more widely acknowledged that secondary BAs are immunomodulatory metabolites that play a part in the regulation of both the innate and adaptive immune systems. Secondary BAs have been shown to decrease IL-1β, IL-6, and TNF-α release in chronic cholestasis, therefore attenuating macrophage-mediated inflammatory responses [32]. There is growing evidence that secondary BAs have a significant impact on adaptive immune responses in addition to their impact on innate immunity. It has been shown that they boost Treg counts, inhibit the development of pro-inflammatory Th17 cells, and reduce the amount of IL-12 generated by dendritic cells. [33,34]. Our serum cytokine analysis demonstrated that DCA significantly decreased the levels of the osteoclastogenic cytokines TNF-α, IL-6, and IL-17 (which is produced by Th17 cells) in the blood. Conversely, DCA increased the levels of anti-osteoclastogenic cytokines such as IL-4 (Th2-associated) and IL-10 (produced from Bregs and Tregs). These findings collectively offer the first concrete proof that DCA has osteo-protective effects by influencing immune cell populations and cytokine profiles that control bone remodelling in addition to inhibiting osteoclastogenesis. DCA protects against LPS-induced inflammatory bone loss *in vivo* by restoring the balance between pro- and anti-osteoclastogenic immune pathways through its targeting of the pro- and anti-inflammatory cells. These results show that DCA has a distinct mode of action that involves both skeletal and immunological regulation, making it a promising therapeutic agent for the treatment of inflammatory bone loss disorders. One drawback of our research is that we have not thoroughly examined how DCA affects osteoblasts’ levels of mRNA expression. Although we carried out a partial analysis, a thorough assessment was not carried out. On the other hand, a more extensive investigation was conducted into the impact of DCA on osteoclast mRNA expression. This gap restricts our comprehension of DCA’s full molecular significance in osteoblast activity.

## Acknowledgment

SY and RKS acknowledge the Department of Biotechnology, AIIMS, New Delhi, India, for providing infrastructural facilities. SY, thank DBT for the research fellowship.

## Conflict of Interest Statement

The authors declare no conflicts of interest.

## Data availability

All data generated or analysed during this study are included in this published article and its supplementary information files. The current study’s data are available from the corresponding author upon reasonable request.

## Funding

This work was financially supported by projects: Department of Science and Technology-Science and Engineering Research Board (EMR/2016/007158), Department of Biotechnology (BT/PR41958/MED/97/524/2021), ICMR (61/05/2022-IMM/BMS), ICMR (EMDR/IG/13/2024-01-00842), sanctioned to RKS.

## Credit authorship contribution statement

RKS contributed to the conception and design of the study; SY contributed to the acquisition and analysis of data; SY contributed to drafting the text or preparing the figures. SR helped with Western blotting, immunofluorescence and helped with drafting. CS did qPCR, and MS helped during *in vivo* studies. PKM provided valuable inputs. All authors reviewed the manuscript. All authors contributed to the article and approved the submitted version.

Send correspondence to:

Dr. Rupesh K. Srivastava,

Additional Professor,

*Translational Immunology, Osteoimmunology & Immunoporosis Lab (TIOIL)*

*An ICMR Collaborating Centre of Excellence on Bone Health*

Department of Biotechnology, All India Institute of Medical Sciences (AIIMS),

New Delhi-110029, India

Cell: +91-9179567399

## Notes

### Competing Interest Statement

The authors have declared no competing interest.

